# Association of 4 epigenetic clocks with measures of functional health, cognition, and all-cause mortality in The Irish Longitudinal Study on Ageing (TILDA)

**DOI:** 10.1101/2020.04.27.063164

**Authors:** Cathal McCrory, Giovanni Fiorito, Belinda Hernandez, Silvia Polidoro, Aisling M. O’Halloran, Ann Hever, Cliona Ni Cheallaigh, Ake T. Lu, Steve Horvath, Paolo Vineis, Rose Anne Kenny

**Affiliations:** Department of Medical Gerontology, Trinity College Dublin, Ireland; Department of Biomedical Sciences, University of Sassari, Italy; Italian Institute for Genomic Medicine, Italy; Department of Clinical Medicine, St. James Hospital, Dublin, Ireland; Department of Human Genetics, David Geffen School of Medicine, Department of Biostatistics Fielding School of Public Health, University of California Los Angeles, USA; MRC Centre for Environment and Health, Imperial College London, UK

**Keywords:** epigenetic clocks, DNA methylation, health, cognition, all-cause mortality

## Abstract

The aging process is characterized by the presence of high interindividual variation between individuals of the same chronical age prompting a search for biomarkers that capture this heterogeneity. The present study examines the associations of four epigenetic clocks - Horvath, Hannum, PhenoAge, GrimAge - with a wide range of clinical phenotypes, and with all-cause mortality at up to 10-year follow-up in a sample of 490 participants in the Irish Longitudinal Study on Ageing. Results indicate that the GrimAge clock represents a step-improvement in the predictive utility of the epigenetic clocks for identifying age-related decline in an array of clinical phenotypes.

## MAIN

Individuals of the same calendar age exhibit marked divergence in their biological aging rate, indicating that the progression of time represents an incomplete account of the aging process ^1^. This simple observation has spawned a search for biomarkers that are sensitive to individual differences in the pace of ageing. A number of candidate biomarkers have been mooted ^2^, but among these, the epigenetic clocks hold arguably the most promise as a sensitive biomarker of ageing. The development of the epigenetic clocks followed from the observation that patterns of DNA methylation - which refers to the addition or removal of methyl (-CH3) groups to Cytosine-phospho-Guanine (CpG) sites - change with age. The advent of DNA methylation microarray technology enabled the identification of the specific genomic locations of CpG sites that were differentially methylated according to age. The most commonly used of what is now referred to as the first-generation clocks are Hannum’s ^3^ blood-specific clock, and Horvath’s ^4^ pan-tissue clock, which are based on levels of DNA methylation at 71 and 353 CpG sites respectively. These clocks are highly correlated with chronological age (r=0.95 in samples with a full age range) ^5^, and the discrepancy resulting from the regression of DNA methylation age on calendar age - a putative measure of biological age acceleration - is associated with increased risk of all-cause mortality ^6,7^. Hereinafter we refer to the age acceleration (AA) estimates derived from the epigenetic clocks with the suffix AA.

In a wide-ranging review describing the development and evolution of the epigenetic clocks, Horvath & Raj ^5^ acknowledged that the first-generation clocks exhibited only weak associations with clinical measures of physiological dysregulation such as blood pressure or glucose metabolism that anticipate hard disease end-points. Recently, Zhang et al. ^8^ have shown that, in principle, a near perfect predictor of calendar age can be estimated from DNA methylation levels, but its association with mortality attenuates with increased accuracy in predicting calendar age; the logical corollary of which is that the training sets being used for deriving accurate biological age estimators require more for their calibration than simply calendar age. The second-generation clocks, PhenoAge ^9^ and GrimAge ^10^ were designed to overcome these limitations by including DNAm correlates of morbidity and mortality. The PhenoAge clock was developed using a two-stage approach. In the first stage, a weighted composite of 10 clinical characteristics (calendar age, albumin, creatinine, glucose, c-reactive protein, lymphocyte percentage, mean cell volume, red blood cell distribution weight, alkaline phosphatase and white blood cell count) was used to develop a phenotypic age estimator. In the second stage, the phenotypic age estimator was regressed on DNAm levels resulting in the identification of 513 CpG sites that exhibited marked differences in disease and mortality among individuals of the same calendar age. Initial validation work by Levine and colleagues confirmed that PhenoAgeAA outperformed the first generation-clocks in the prediction of many age-related diseases and lifespan. Lu and colleagues ^10^ adopted a slightly different strategy in the development of the latest clock – GrimAge. In the first stage, they identified DNAm based surrogates of 12 plasma proteins as well as smoking pack years. In the second stage, they regressed time-to-death due to all-cause mortality on the DNA methylation based-markers of plasma protein levels and smoking pack-years, identifying 1030 CpG sites that jointly predicted mortality risk.

Given the novelty of these measures the validation of the second-generation clocks has only really begun in earnest, but early studies suggest that they represent an important step forward in our ability to predict a range of important age-related health outcomes and lifespan. While not the primary focus of the paper, McCrory et al.^11^ reported that PhenoAgeAA was associated with slower performance on the timed-up-and-go (TUG) task, and with higher risk of prefrailty/frailty, but not with activity limitations in a paper comparing the performance of allostatic load and the epigenetic clocks for advancing our understanding of how low socio-economic position gets biologically embedded. A separate paper by the same group ^12^ noted that PhenoAgeAA correlated 0.21 with allostatic load – a compositive measure of physiological dysregulation – while the correlation of HorvathAA and HannumAA with AL was close to zero in the overall sample, and higher for men compared with women.

Maddock et al. ^13^ examined associations of HorvathAA, HannumAA, PhenoAgeAA and GrimAgeAA with 3 measures of functional health (grip strength, chair-rise speed, and lung function) and two cognitive performance measures (word recall and mental speed) between the ages of 45-87 using data for 3 British cohorts: the 1946 National Survey of Health and Development (NSHD), 1958 National Child Development Study (NCDS), and Twins UK. They found that the first-generation clocks were not significantly associated with any of the 5 outcome measures in the meta-analysis of these cohorts, but that the second-generation clocks were. Specifically, PhenoAgeAA was associated with lower grip strength, worse lung function, and slower mental speed; while GrimAgeAA was associated with worse lung function, poorer word recall, and slower mental speed. Importantly, GrimAgeAA as assessed at baseline was associated with a faster rate of decline in grip strength and lung function between 53-69 years of age in the NSHD cohort.

In one of the most comprehensive investigations to date, Hillary & collaborators ^14^ present further evidence that the GrimAge clock exhibits remarkable promise as a biomarker of aging. Using data for some 709 individuals (mean age=73) participating in the 1936 Lothian Birth Cohort (LBC), they reported that GrimAgeAA was inversely associated with lung function, levels of iron, LDL and total cholesterol; and positively associated with measured weight, body mass index, C-reactive protein, creatinine, and interleukin 6. GrimAgeAA was also found to be negatively associated with performance on a battery of cognitive ability measures including, choice reaction time, digit symbol-coding, symbol search, matrix reasoning, and general intelligence, decreased brain region volume, and increased white matter hyperintensities. It also predicted increased risk of all-cause mortality with an 81% increase in hazard per standard deviation increase in GrimAgeAA.

While the early findings do indeed seem to indicate that the second-generation clocks may be capturing important aspects of accelerated biological ageing, Dugue ^15^ in a recent critique of the epigenetic clocks cautioned that early studies generally report stronger associations than later studies and are more likely to be affected by publication bias. This study represents further evaluation of the new generation of clocks using a sub-sample of 490 participants to the Irish Longitudinal Study of Ageing (TILDA), a nationally representative study of community-dwelling older people in Ireland. A unique feature of TILDA in the context of the wider family of ageing studies worldwide is that all participants are invited to undergo a comprehensive clinical health assessment, including the collection of blood samples, so it has much deeper phenotyping of individuals than is typical in an epidemiological study. The assessment takes approximately 3 hours to complete and provides data for a comprehensive set of health and cognitive domains that are relevant to ageing. This novel study provides important new evidence concerning the relevance of the epigenetic clocks for clinical medicine by examining the utility of 4 epigenetic age acceleration indices – HorvathAA, HannumAA, PhenoAgeAA, and GrimAgeAA – for predicting functional decline, disability, co-morbidity, and cognitive impairment / dementia using a series of measures that are well validated in the clinical literature as markers of age-related decline including walking speed, grip strength, Fried frailty, polypharmacy, the Montreal Cognitive Assessment (MOCA) and the Mini-Mental State Exam (MMSE). We also examine the utility of epigenetic AA for predicting time to event due to all-cause mortality at up to 10-year follow-up.

## RESULTS

Figure 1 depicts the functional form of change in each of the dependent variables according to age and sex using the full TILDA sample, demonstrating clearly that our various outcome measures are highly sensitive to age-related decline. It shows that walking speed and grip strength decline steadily with age, and that Fried frailty scores increase with age, as does the prevalence of polypharmacy. Likewise, the number of errors on the Montreal Cognitive Assessment, Mini-Mental State Exam (MMSE) and Sustained Attention Reaction Time task increases sharply with age, as does performance on the Choice Reaction Time (CRT) task. Normative values and age gradients for many of these measures derived using the TILDA dataset have been published elsewhere ^16^. Table 1 describes the characteristics of the sample and reports mean scores (SD) as well as the range for each of the functional health and cognitive performance measures. The mean age of the sample was 62.2 and 50.2% were female. The magnitude of the Pearson intercorrelations between HorvathAA, HannumAA, and PhenoAgeAA for the TILDA epigenetic sample have been previously reported in ^12^ so we report only the correlations of GrimAgeAA with the other 3 epigenetic AA measures. The correlation of GrimAgeAA with HorvathAA (r = −0.02) and HannumAA (r = 0.00) was negligible, although it was modestly correlated with PhenoAgeAA (r = 0.40).

**Figure 1.**
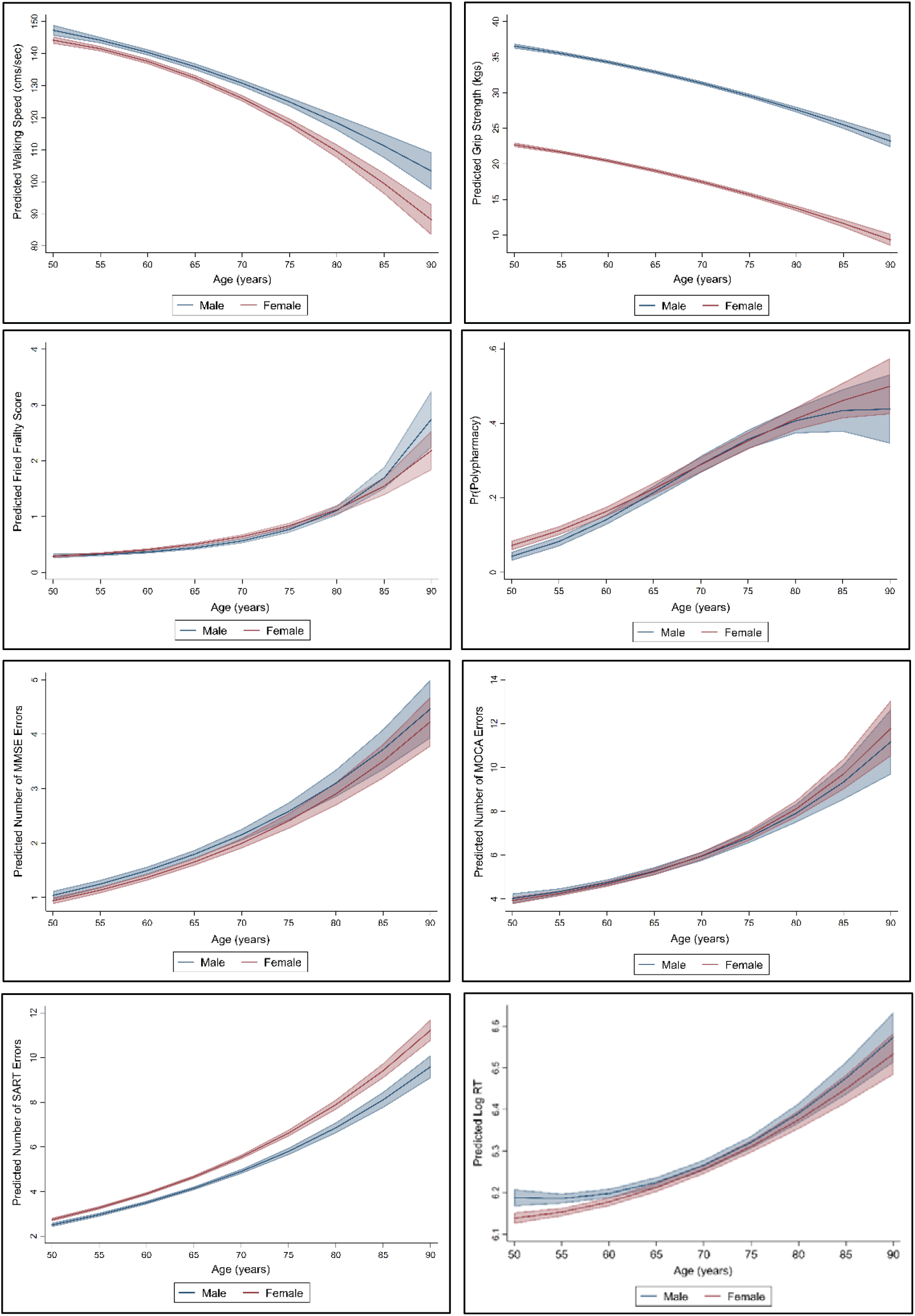
Predicted values for various indices of physical and cognitive functioning according to age and sex in the Irish Longitudinal Study of Ageing (TILDA)

**Table 1:** Baseline characteristics of the TILDA epigenetic sub-sample.

|  | Mean (SD) or n (%) |
| --- | --- |
| HorvathAA | 0.00 (7.4) |
| HannumAA | 0.00 (7.3) |
| PhenoAgeAA | 0.00 (5.0) |
| GrimAgeAA | 0.00 (4.3) |
| Age | 62.2 (8.3) |
| Female Sex | 246 (50.2%) |
| Social class trajectory |  |
| - Stable high | 123 (25.1%) |
| - Upwardly mobile | 125 (25.5%) |
| - Downwardly mobile | 121 (24.7%) |
| - Stable low | 121 (24.7%) |
| Smoking |  |
| - Never smoked | 193 (39.4) |
| - Past smoker | 211 (43.1) |
| - Current smoker | 86 (17.6) |
| Hazardous drinking |  |
| - No | 371 (75.7%) |
| - Yes | 77 (15.7%) |
| - Missing | 42 (8.6%) |
| Physical Activity |  |
| - Low | 142 (29.0%) |
| - Moderate | 175 (35.7%) |
| - High | 167 (34.1%) |
| - Missing | 6 (1.2%) |
| BMI | 28.8 (5.1) |
| Walking Speed (cm/sec) | 136.1 (21.6) |
| Grip strength (kgs) | 26.5 (9.4) |
| Polypharmacy | 94 (19.2%) |
| Fried Frailty score | 0.43 (0.74) |
| MOCA errors | 4.9 (3.4) |
| MMSE errors | 1.5 (1.9) |
| SART errors | 3.7 (4.0) |
| Raw CRT (ms) | 505.7 (139.2) |

Figure 2 shows the change in each of the clinical health and cognitive performance measures associated with a one-year increase in epigenetic AA according to each of the four clocks while adjusting for calendar age and sex in the minimally adjusted models (model 1), and additionally for life course socio-economic characteristics, smoking, hazardous levels of alcohol consumption, physical activity levels, and body mass index (BMI) in the fully adjusted models (model 2). The unstandardized beta coefficients (B), odds ratios (OR), and incident rate ratios (IRR) and associated 95% confidence intervals are reported in Table 2. There was only one significant association of the first-generation clocks with any of the outcome measures under investigation. HorvathAA was associated with higher grip strength in both the minimally and full multivariable adjusted models.

**Figure 2.**
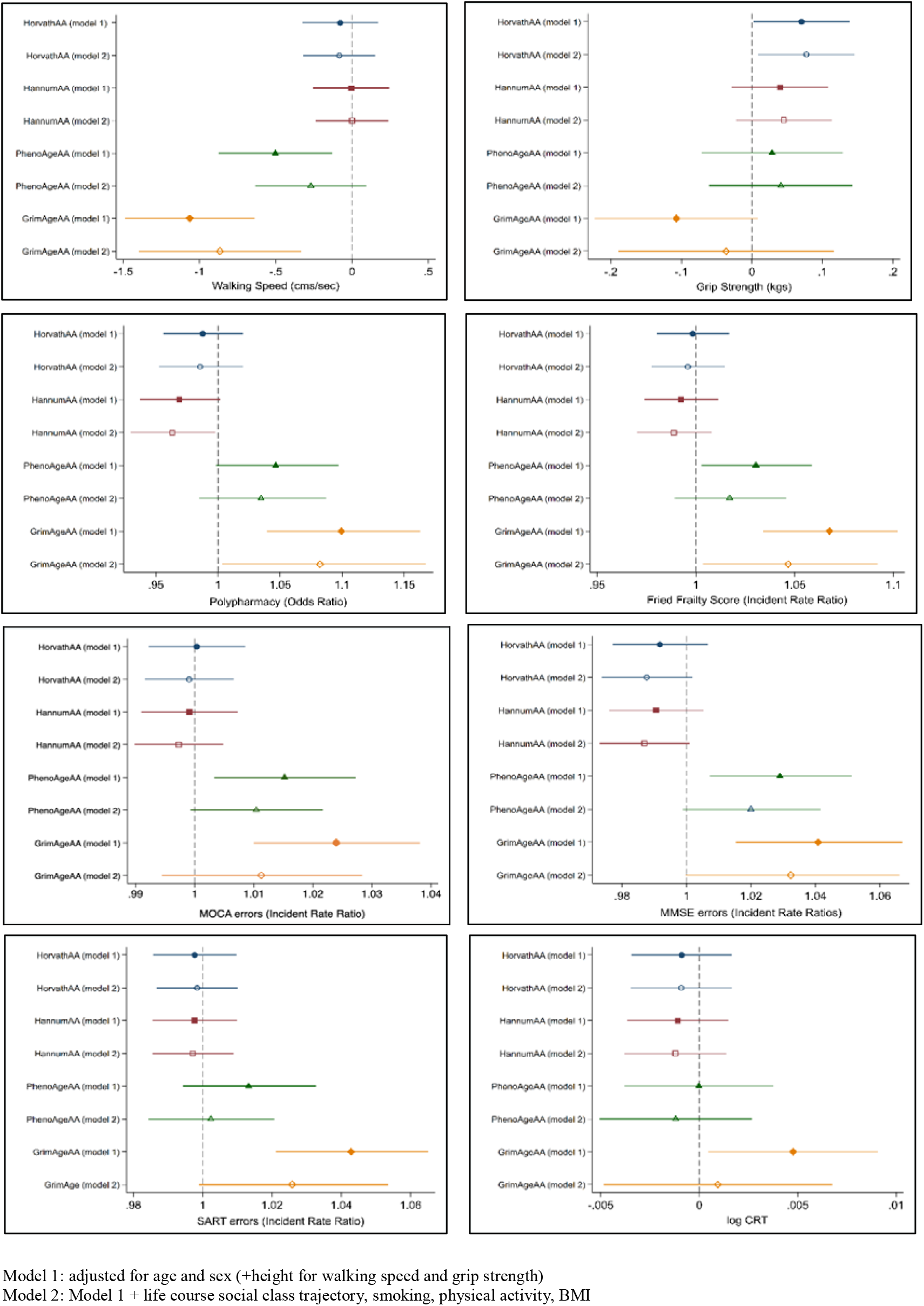
Association of cell intrinsic epigenetic age acceleration measures with physical and cognitive functioning in the minimally (model 1) and full multivariable adjusted models (model 2)

**Table 2:**
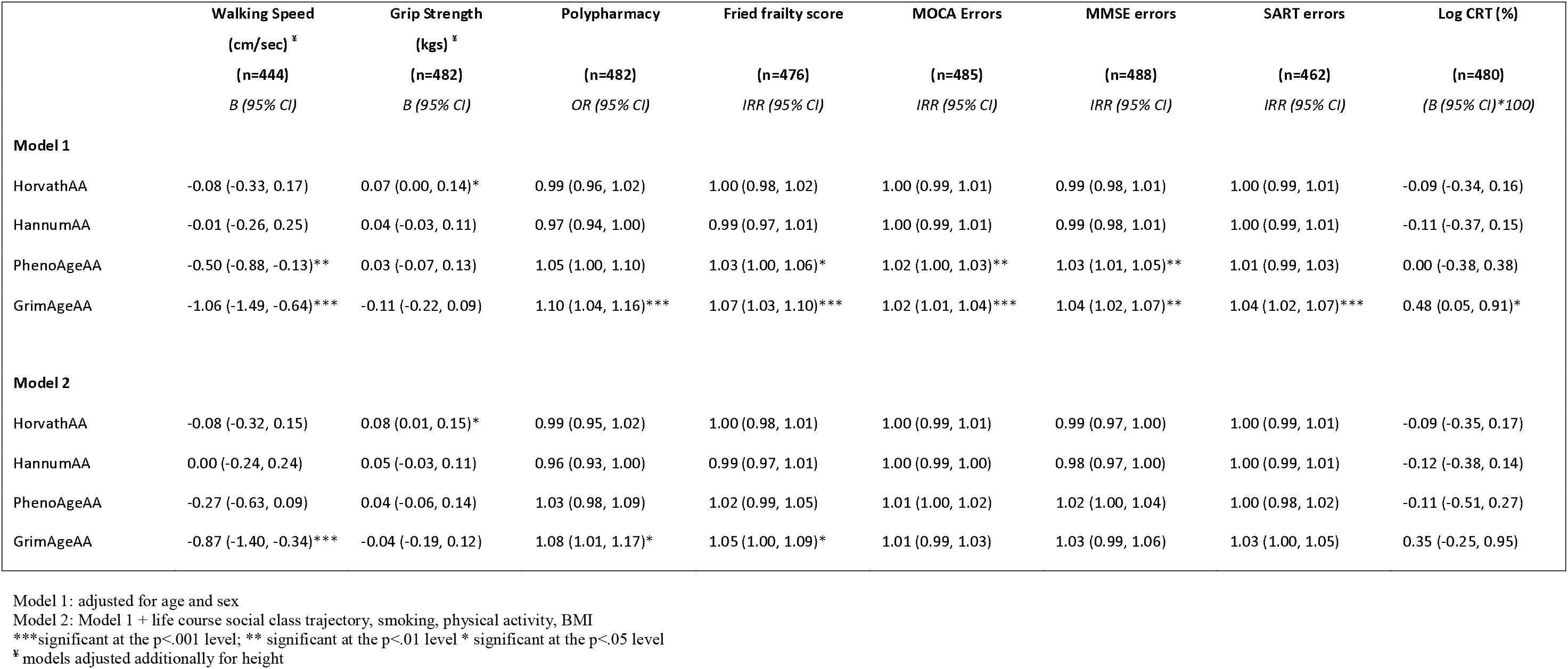
Association of cell intrinsic epigenetic age acceleration measures with physical and cognitive functioning in the minimally (model 1) and full multivariable adjusted models (model 2)

PhenoAgeAA was associated with 4/8 physical and cognitive outcomes in the minimally adjusted models including, slower walking speed (B= −0.50, 95% CI = −0.88, −0.13; p=0.009), higher Fried frailty (IRR = 1.03, 95% CI = 1.00, 1.06; p=0.030), MOCA errors (IRR = 1.02, 95% CI = 1.01, 1.04; p=0.012), and MMSE errors (IRR =1.03, 95% CI = 1.01, 1.05; p=0.009), but none of these associations survived multivariable adjustment. GrimAgeAA was associated with 7/8 physical and cognitive health outcomes in the minimally adjusted models: slower walking speed (B= −1.06, 95% CI = −1.49, −0.64; p<0.001), increased polypharmacy (OR = 1.10, 95% CI = 1.04, 1.16; p<0.001), higher Fried frailty score (IRR = 1.07, 95% CI = 1.03, 1.10; p<0.001), MOCA errors (IRR = 1.02, 95% CI = 1.01, 1.04; p<0.001), MMSE errors (IRR =1.04, 95% CI = 1.02, 1.07; p=0.002), SART errors (IRR=1.04, 95% CI = 1.02, 1.07; p<0.001), log CRT*100 (B=0.048, 95% CI = 0.05, 0.91; p=0.030); and continued to be associated with 3/8 outcomes (walking speed, frailty score and polypharmacy) in the full multivariable adjusted models.

Table 3 (model 1) shows that a one year increase in GrimAgeAA was associated with a 18% increase (95% CI = 1.09, 1.28; p<0.001) in the hazard of all-cause mortality at up to 10-year follow-up, and these estimates were only marginally affected when adjusted for socio-economic characteristics and lifestyle factors (HR=1.16, 95% CI=1.05, 1.29; p=0.004) (Table 1, model 2). None of the other epigenetic AA measures significantly predicted mortality.

**Table 3:** Hazard Ratio for all-cause mortality at up to 10-year follow for each of the four epigenetic AA measures in the minimally and full multivariable adjusted models (n=489).

|  | **Model 1** | **Model 2** |
| --- | --- | --- |
|  | HR (95% CI) | HR (95% CI) |
| HorvathAA | 1.00 (0.95, 1.04) | 1.00 (0.96, 1.05) |
| HannumAA | 0.98 (0.94, 1.03) | 0.99 (0.95, 1.03) |
| PhenoAgeAA | 1.05 (0.98, 1.12) | 1.02 (0.96, 1.09) |
| GrimAgeAA | 1.18 (1.09, 1.28)*** | 1.16 (1.05, 1.29)** |

\*\*\*significant at the p<.001 level; \*\*significant at the 0.01 level

### Sensitivity check

As described in the methods section, we calculated the so called intrinsic epigenetic AA for use in the analysis. As a sensitivity check on our results, we also estimated the relationship between cell extrinsic epigenetic AA (i.e. adjusted for calendar age but not white blood cell percentages) and the various outcomes measures under investigation (Supplementary Table 1). In general, results were consistent with what we found for cell intrinsic epigenetic AA so here we discuss only the differences. PhenoAgeAA was now additionally associated with polypharmacy (OR= 1.06, 95% CI = 1.01, 1.11; p=0.024), and GrimAgeAA was now additionally associated with lower grip strength (B= −0.14, 95% CI = −0.25, −0.02; p=0.019) in the minimally adjusted, but not the full multivariable adjusted models. Adjusting for socio-economic characteristics and other lifestyle factors, PhenoAgeAA was associated with 1/8 physical health and cognitive outcomes (i.e. MMSE errors) while GrimAgeAA remained associated with 4/8 outcomes (i.e. slower walking speed, polypharmacy, Fried frailty score, MOCA errors).

## DISCUSSION

This study compared the utility of the so-called first and second-generation epigenetic clocks for predicting functional health, cognitive / neuropsychological performance, and all-cause mortality in a sub-sample of TILDA participants confirming that the new GrimAge clock represents a strong marker of premature ageing with huge implications for clinical medicine. The results of this study confirm Horvath and Raj’s ^5^ previous observation that the first-generation clocks are not sensitive predictors of age-related decline in clinical health measures. Few of the associations were significant, and even among those that were, the observed relationships were in the opposite direction to what one would hypothesise. A one-year increase in HorvathAA was paradoxically associated with higher grip strength, which may be a chance finding given the number of comparisons performed. The mostly null associations for the first-generation clocks are consistent with the results obtained by Maddock et al. ^13^ in a recent study involving the British cohort studies.

The second-generation clocks by contrast markedly outperformed the first-generation clocks in the prediction of many functional health and cognitive performance measures, and GrimAgeAA estimated risks were about twice those estimated for PhenoAgeAA. PhenoAgeAA was associated with slower walking speed, higher Fried frailty, MOCA errors and MMSE errors, but these relationships no longer held when adjusted for socio-economic characteristics and other lifestyle-related factors. GrimAgeAA by contrast was associated with all outcome variables under investigation, except grip strength, and continued to be associated with four of them (walking speed, polypharmacy, Fried frailty score, and all-cause mortality) even in the full multivariable adjusted models; a finding which implies that the GrimAge clock is tapping variation in the pace of ageing that is not simply due to socio-economic position (SEP) and other lifestyle-related factors. This is despite the fact that GrimAgeAA is more strongly associated with SEP than the other 3 clocks. McCrory et al. ^11^ have previously shown that HorvathAA, HannumAA, and PhenoAgeAA are not strongly related to socio-economic characteristics in the TILDA epigenetic sample, but in this study, having a stable low SEP across the life course (i.e. low childhood social class / low adulthood social class) was associated with 2.42 years (95% CI = 1.36, 3.49; p<0.001) of epigenetic AA according to the GrimAge clock compared with those who were stable high (i.e. high childhood social class / high adulthood social class) in SEP. Being a current smoker was associated with 7.91 years (95% CI = 7.06, 8.76; p<.001) of GrimAgeAA, being a hazardous drinker according to the CAGE questionnaire was associated with 0.94 years (−0.13, 2.02, p=0.086) of GrimAgeAA, while levels of physical activity were inversely associated with GrimAgeAA. Finally, BMI was found to be unrelated to GrimAgeAA (Figure 3).

**Figure 3:**
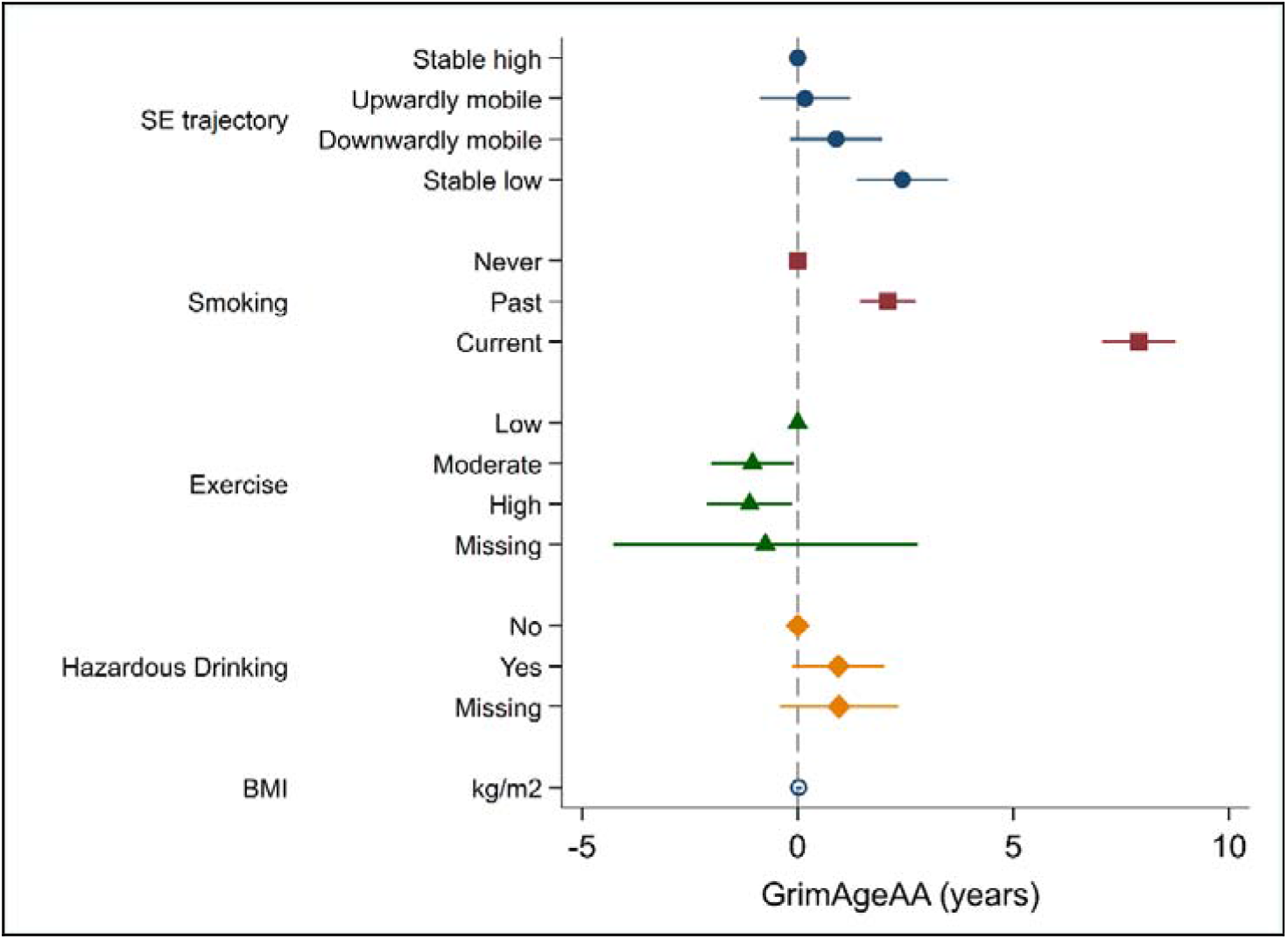
Age and sex adjusted independent association of cell intrinsic GrimAgeAA with socio-economic characteristics and other lifestyle factors.

Other factors which we did not adjust for in this analysis but may mediate the association of GrimAgeAA with our broad array of clinical outcomes includes: (i) genetics, as the GrimAge clock was calibrated by regressing time to death due to all-cause mortality on DNA methylation levels; (ii) life course psychosocial stressors, which have been linked with accelerated epigenetic ageing ^17^ and physiological wear and tear in tissues and organ systems ^18^; (iii) nutritional deficiencies, particularly folate insufficiency, which is a key nutritional factor in one-carbon metabolism ^19^; and (iv) exposure to environmental toxins ^20^, which may elicit epigenetic modifications ^21^, and are known to be deleterious to health. Future research should be designed to address these possibilities.

Shiels and colleagues ^22^ contend that a sequitur for any valid biomarker of ageing is that it shows a statistically significant association with a measure of health or organ functional capacity over and above calendar age. We went a step further to examine whether epigenetic AA measures are *clinically* relevant as well as statistically relevant. On the basis of previous exploratory work involving HorvathAA, HannumAA and PhenoAgeAA, McCrory and collaborators ^12^ questioned whether the residual from the regression of DNA methylation age on chronological age represented the sine qua non of biological ageing. The new GrimAge clock seems to allay many of these concerns, reaffirming the belief that DNA methylation levels at very few CpG sites across the human methylome may be informative about physical health and brain health independently of calendar age. Moreover, the GrimAge clock is highly stable over time with a mean intraclass correlation coefficient of 0.85 across 4 measurement occasions spanning approximately a decade, which is clearly a desirable feature to have in a biological age predictor ^14^.

Nevertheless, what remains to be established is whether epigenetic AA predicts changes in health and cognition prospectively, and perhaps more pertinently, whether change in the epigenetic AA measures is associated with change in physical health and cognitive performance. This is important in order to help establish whether DNA methylation is a cause or a consequence of aging. To date the evidence is inconclusive. For example, the study involving the LBC found that GrimAgeAA was associated with a borderline significant decline in cognition over time that was attenuated after adjustment for age 11 cognitive ability. Maddock et al. ^13^ found that GrimAgeAA was associated with a faster rate of decline in grip strength and lung function, and PhenoAgeAA with faster decline in lung function between 53 and 69 years of age in the NSHD cohort. Change in the AA measures was available for a sub-sample of NSHD participants but there was little evidence to support the proposition that change in AA was associated with change in the physical and cognitive performance measures. Unfortunately to date, few studies have measured change in both health and patterns of DNA methylation over time, even though these studies are urgently needed to help disambiguate the temporal ordering of the observed relationships. Levine ^23^ recently suggested that the reason the second-generation clocks outperform their progenitors is because they were trained on longitudinal data. Although speculative, it is highly likely that the 3rd generation clocks will be trained on data that assesses changes in DNA methylation levels and change in the type of age-related phenotypes measured so comprehensively in the TILDA study prospectively. There is also a concomitant need to explore the functional significance of genes that are regulated by the CpG sites that are most informative of biological ageing, as this may will inform efforts to compress disease and morbidity and advance precision medicine.

The study benefits from the use of a deeply phenotyped community-dwelling older cohort for whom we have a rich array of clinical health and cognitive performance measures administered in a health centre conducted by trained nurses using standardised operating protocols. We also capture older, frailer individuals as a nurse also administers a home health assessment to those who cannot attend the center and who may be lost to follow-up in other studies. Compared with Maddock et al., ^13^ our study assesses a larger number of health and cognitive outcomes, as well as all-cause mortality. Compared with Hillary’s study ^14^, our study reports results for 4 epigenetic AA measures as opposed to just GrimAgeAA, which allows us to assess using the same pool of participants how the performance of the epigenetic clocks has evolved over time, showing this step change in predictive utility with each iteration. Our study also has a number of weaknesses, perhaps the most notable of which is the relatively small (by epidemiological standards) sample size, and the selective nature of the sample which was originally designed to look at the impact of life course socio-economic trajectories on epigenetic ageing rates.

This study provides important new evidence concerning the utility of the new GrimAge epigenetic clock for predicting age-related decline in a number of important health and cognitive domains relevant to clinical medicine. The fact that GrimAgeAA was so strongly associated with health-span and lifespan is potentially exciting as it holds out the tantalising prospect that we may have identified a biological ageing surrogate with strong theoretical underpinnings that has prognostic utility as an indicator of a person’s state of health. Recent work suggests that epigenetic ageing is potentially reversible ^24^ so the epigenetic clocks may also represent a means for quantifying the efficacy of interventions designed to retard or reverse the ageing process – the holy grail of Geroscience. In summary, evidence is accumulating that DNA methylation plays a central role in the ageing process, and the epigenetic clocks may represent a highly accurate means for quantifying the rate of biological decay in what has been heralded the new era of precision medicine.

## ONLINE METHODS

### Ethics Statement

Ethical approval for the TILDA study was obtained from the Trinity College Dublin Research Ethics Committee and signed informed consent was obtained from all participants.

### Participants

The TILDA study design and sample selection is described in detail elsewhere 1,2. Briefly, the study involves a nationally representative sample of 8,175 community dwelling older persons aged 50 years and over at baseline. Respondents complete a 90-minute computer-assisted personal interview (CAPI) in the home and a separate self-completion questionnaire containing more sensitive information that is returned via post. All participating respondents are invited to attend for a detailed clinic-based health assessment administered by trained nursing staff in either a center-based or home-based setting. The present study uses a sub-sample (n=490) of the baseline TILDA cohort who were selected into the epigenetic study based on their life course social class trajectory as described in 3.

### Epigenetic Age Acceleration (AA) Measures

DNAm levels were assessed using the Infinium Human Methylation 850k Beadchip (Illumina Inc. San Diego, CA). HorvathAA, HannumAA, PhenoAgeAA and GrimAgeAA were estimated from whole blood according to procedures described in Fiorito et al.^4^ and Lu et al.^5^. Since epigenetic AA could be correlated with chronological age and white blood cell (WBC) percentage, we utilised the so-called ‘intrinsic’ epigenetic AA 6 in the analysis, defined as the residuals from the linear regression of epigenetic AA with chronological age and WBC percentages. The latter were estimated using the Houseman et al. ^7^ algorithm. By definition, these measures of AA exhibit zero correlation with calendar age and WBC percentage.

### TILDA Health Assessment

TILDA participants were invited to undergo a detailed health assessment described in 8. If a respondent could not attend the health center but was agreeable to completing a health assessment, a trained nurse administered a subset of the tests in the respondent’s home. Walking speed was only measured in the health center but all other measures were administered at both center-based and home-based assessments.

### Physical Health Outcome Measures

#### Walking Speed

Walking speed was assessed using a 4.88 metre (m) computerized walkway with embedded pressure sensors (GAITRite, CIR Systems Inc., New York, NY). Participants completed two walks along the mat at their normal walking speed. Each trial started 2.5 m before and ended 2 m after the walkway in order to allow room for acceleration and deceleration. The average of the two readings represented the overall walking speed measure expressed in centimetres travelled per second (cm/sec).

#### Grip strength

Grip strength was measured using a Baseline Hydraulic Hand Dynamometer, which consisted of a gripping handle with a strain gauge and an analogue reading scale that increased in 2-kilogram (kg) increments. Respondents were instructed to hold the device in their hand with the forearm at a right angle to their upper arm. The participant was then instructed to squeeze as hard as they could for a few seconds. Results were rounded to the nearest whole number (i.e., if the dial fell between 22 and 24 kg, grip strength was recorded as 23 kg). This procedure was repeated twice in each of the dominant and non-dominant hands and the average of the 4 measurements was used to indicate grip strength.

#### Polypharmacy

Doctor prescribed medication usage was captured as part of the household interview and confirmed by cross-checking the labels on the medicinal packaging. The international non-proprietary name (INN) for any regularly taken medications was assigned and coded using Anatomic Therapeutic Classification (ATC) codes. Polypharmacy was defined as currently taking 5+ doctor prescribed medications and used as a binary variable in the analysis. Previous studies have documented strong concordance between self-reported interview data and pharmacy records ^9,10^.

#### Fried Frailty Score

Frailty was operationalised using TILDA population-specific cut-points following the methodology of Fried and colleagues.^11,12^. Weakness: Mean grip-strength was calculated from an average of two measurements on the dominant hand using baseline dynamometer. Weight and height were measured using standardised procedures during the health centre assessments. Participants scoring in the 20th percentile of sex- and BMI-adjusted grip-strength were given a score of 1, those above the 20th percentile scored 0. Physical activity: Kilocalories (kcals) of energy expended were calculated from the International Physical Activity Questionnaire-Short Form [IPAQ-SF]). Participants scoring in the 20th percentile of sex-adjusted kcals were given a score of 1, those above the 20th percentile scored 0. Slow walking speed: Slow walking speed: recorded time taken in seconds (s) to perform the Timed Up-and-Go (TUG) task. Participants scoring in the slowest 20th percentile of sex- and height-adjusted TUG time were given a score of 1, those above the slowest 20th percentile scored 0. Unintended Weight loss: was ascertained by the question ‘In the past year have you lost 10 pounds (4.5 kg) or more in weight when you were not trying to’. Participants who responded ‘yes’ were given a score of 1, those who responded ‘no’ scored 0. Exhaustion: was captured using two items from the 20-item Centre for Epidemiological Studies Depression (CES-D) scale. Participants were asked how often they felt that ‘I could not get going’ and ‘I felt that everything I did was an effort’. Participants who responded ‘a moderate amount/all of the time’ to either question were given a score of 1, those who responded ‘none/some of the time’ to both questions scored 0. The total score (range 0-5) was modelled as the response variable.

### Global Cognitive / Neuropsychological Outcome Measures

#### Mini-Mental State Exam (MMSE)

The MMSE ^13^ was originally designed to provide a brief, standardized assessment of mental status in psychiatric patients, although it is now used widely in clinical and epidemiological settings as a screening tool for the assessment of global cognitive impairment/dementia. It takes approximately 5-10 minutes to administer and assesses attention and concentration, memory, language, visuo-constructional skills, calculation and orientation. Because the instrument suffers from ceiling effects, we used the count of errors in the analysis (i.e. 30 – number of items incorrect) as the dependent variable.

#### Montreal Cognitive Assessment (MOCA)

Global cognitive status was also measured using the Montreal Cognitive Assessment (MOCA) ^14^. The MOCA was designed as a brief cognitive screening tool to detect mild cognitive impairment (MCI) among community and clinic-based samples. The test takes approximately 10 minutes to administer and assesses performance across a number of cognitive domains including: executive functioning/visuospatial abilities, attention/concentration, language; animal naming, short term memory, abstraction, and orientation. The instrument yields a sum score ranging from 0-30. Recent studies involving clinical samples suggest the MOCA performs better than the MMSE in detecting early signs of cognitive decline 15 and hence servers as a useful epidemiological screening tool for identifying the transitional state between normal cognitive functioning and more severe cognitive impairment. Again, we modelled the count of errors as the dependent variable (i.e. 30 – number of items incorrect).

#### Sustained Attention Reaction Time Task (SART)

The ability to sustain attention is an important component of normal executive functioning, and normal cognitive ageing is associated with declines in selective (i.e. ability to attend) and divided (i.e. ability to focus on multiple tasks simultaneously) attention ^16^. The SART is a computerised performance reaction-time (RT) task. It requires participants to attend to a repeating stream of digits (1-9) and to press the response key for each number that they see (GO trials), except number ‘3’ (NO GO trials). Each digit from 1 through 9 appears for 300 milliseconds (ms), with an interval of 800ms between digits, and the cycle is repeated 23 times giving a total of 207 trials. The task takes approximately 4 minutes to complete. Commission errors (i.e. responding to NO–GO trials) reflect lapses of sustained attention, and omission errors (i.e. failure to respond to GO trials) reflect a break from task engagement, also corresponding to lapsing attention. The total count of commission errors made on the task was modelled as the response variable.

#### Choice Reaction Time Task (CRT)

Processing speed slows in ageing irrespective of whether it is measured using psychometric tests (e.g. colour trails), cognitive-experimental tasks (e.g. reaction time) or psychophysical tasks (e.g. inspection time) leading some to speculate that it may represent a biomarker of cognitive ageing ^17^. Participants completed a standard 2-choice reaction time task as part of a detailed neuropsychological battery. They were instructed to rest their finger on a ‘home’ plate and wait for a stimulus to appear on screen stating ‘Yes’ or ‘No’. They then had to move their finger from the home plate to the corresponding ‘Yes’ or ‘No’ button. Participants completed several practice trials in the presence of the clinical nurse to ensure familiarity with the test procedure. Respondents were only permitted to enter the test phase if they demonstrated enough understanding of the requirements of the task during the test phase. The trial phase consisted of a series of 100 trials with the stimulus prompt alternating in a randomized sequence and appearing approximately 50% of the time for each stimulus response. RT can be decomposed into a decision time (interlude between stimulus onset and movement from the home plate) and movement time component (interlude between movement from home plate to the relevant response key). As the CRT data exhibited marked positive skew, it was log transformed prior to analysis.

#### All-Cause Mortality

In total, 35/490 participants or 7.4% of the epigenetic sub-sample were confirmed as deceased at up to 10-year follow-up (i.e. by end of wave 5 sweep of data collection in 20th December 2018). Mortality status was available from two sources: (1) through data linkage to the General Registrar’s Office (GRO) National Death Registry which was done in March 2017; and (2) via an End of Life Interview (EOL) for deaths occurring subsequent to administrative record linkage, which was conducted with the respondent’s next of kin (if available). Date of death was available for 17/35 respondents from the GRO and for a further 7/35 via the EOL interview. Time of death was unavailable for the remaining 11 cases because no EOL was completed or because TILDA allows a period of 6 months to elapse before attempting to conduct an EOL interview with the spouse/family of recently deceased cohort members. In these instances, we imputed date of death as half of the mean between-wave interval after the ‘date of last contact’.

#### Covariates

In addition to age (years) and sex (male, female), we controlled for other socio-economic characteristics and lifestyle factors that could potentially confound the association of the putative biological ageing measures with the various outcome measures. Life course social class trajectory was measured using the cross-classification of participants’ childhood (i.e. father’s) and adulthood (i.e. own contemporaneous) social class into 4 categories as follows: stable professional, upwardly mobile, stable unskilled, and downwardly mobile. Smoking history is represented using a 3-level variable: Never Smoked, Past Smoker, Current Smoker. Physical activity was assessed using the eight-item short form of the International Physical Activity Questionnaire (IPAQ) (24). It measures the amount of time (minutes) spent walking and engaged in moderate and vigorous physical activity, and the amount of time spent sedentary and categorised into: Low, Medium, and High levels of physical activity as per the IPAQ protocol (www.ipaq.ki.se). Hazardous drinking was assessed using the CAGE alcohol screening test ^18^. The scale comprises four items and follows a dichotomous yes/no response format. Answering yes to two or more questions indicates a clinically significant profile and constitutes potentially hazardous drinking. Body Mass Index (BMI) was calculated from measured height and weight. Height was measured using a SECA 240 wall mounted measuring rod and weight was measured using a SECA electronic floor scales.

#### Treatment of missing cases

The overall level of missing data for the covariates was small. Six individuals were missing on the physical activity measure and 42 individuals were missing on the alcohol consumption measure; so, we coded these as “missing” using dummy variables so that they would not be lost to the analysis. One person was missing data for height and weight. The amount of missing data on the outcome measures was small overall and generally due to non-systematic sources of variance (e.g. technical problems).

### Statistical Analysis

All analyses were conducted using Stata 15.0 (StataCorp, College Station, TX, USA). The outcome measures were regressed separately on each of the four epigenetic age acceleration measures adjusting for chronological age (years), and sex using ordinary least squares regression (walking speed, grip strength, reaction time), Poisson regression (frailty scores), negative binomial regression (MOCA errors, MMSE errors, SART commission errors), logistic regression (polypharmacy), and Cox regression (mortality) as appropriate. This shows the change in each of the outcome measures associated with a one-year increase in AA in each of the four biological ageing measures. In the analyses involving walking speed and grip strength, we adjusted additionally for measured height (cms) which differs between men and women and is strongly associated with stride length and strength. In secondary models we adjusted additionally for participants’ life course socio-economic trajectory in order to take account of characteristics associated with selection into the sample, as well as smoking, hazardous alcohol consumption, physical activity levels, and BMI to determine whether any putative association of the age acceleration measures with health survived adjustment for lifestyle factors. We tested for effect modification by sex by fitting separate sex*EAA interaction terms with respect to each outcome in the minimally adjusted models. Only two of the 32 contrasts were significant – both on the SART task, but in opposite directions. Women made a significantly higher number of commission errors on the SART task per one-year increase in GrimAgeAA, but significantly fewer errors per one-year increase in HannumAA. We report results therefore for the overall sample.

## Acknowledgements

This work was supported by the Health Research Board (HRB) of Ireland under an Emerging Investigator Award (EIA-2017-012) to CMC. This work was also supported by the LIFEPATH grant to P.V. at Imperial College London (European Commission H2020 grant, Grant number: 633666). Funding for the TILDA project was supported by the Irish Government, the Atlantic Philanthropies, and Irish Life plc. SH and ATL were supported by NIH/NIA IU01AG060908. The funders had no involvement in the study design, collection, analysis and interpretation of data, or authorship of the submitted work.

