## Supplementary Table 1 for "Association of 4 epigenetic clocks with measures of functional health, cognition, and all-cause mortality in The Irish Longitudinal Study on Ageing (TILDA)"

**Supplementary Table 1: Association of cell extrinsic epigenetic age acceleration measures with various indices of physical and mental functioning in the minimally (model 1) and full multivariable adjusted models (model 2)**

|  | **Walking Speed (cm/sec) ^¥^**  **(n=444)** | **Grip Strength**  **(kgs) ^¥^**  **(n=482)** | **Polypharmacy**  **(n=482)** | **Fried frailty score**  **(n=476)** | **MOCA Errors**  **(n=485)** | **MMSE errors**  **(n=488)** | **SART errors**  **(n=462)** | **Log CRT (%)**  **(n=480)** |
| --- | --- | --- | --- | --- | --- | --- | --- | --- |
|  | *B (95% CI)* | *B (95% CI)* | *OR (95% CI)* | *IRR (95% CI)* | *IRR (95% CI)* | *IRR (95% CI)* | *IRR (95% CI)* | *(B (95% CI)*100)* |
| **Model 1** |  |  |  |  |  |  |  |  |
| HorvathAA | -0.06 (-0.30, 0.19) | 0.08 (0.01, 0.15) | 0.99 (0.95, 1.02) | 1.00 (0.98, 1.01) | 1.00 (0.99, 1.01) | 0.99 (0.98, 1.01) | 1.00 (0.99, 1.01) | -0.09 (-0.36, 0.15) |
| HannumAA | -0.01 (-0.24, 0.26) | 0.05 (-0.02, 0.12) | 0.97 (0.94, 1.00) | 0.99 (0.97, 1.01) | 1.00 (0.99, 1.01) | 0.99 (0.98, 1.01) | 1.00 (0.99, 1.01) | -0.11 (-0.37, 0.14) |
| PhenoAgeAA | -0.53 (-0.89, -0.16)** | 0.01 (-0.09, 0.11) | 1.06 (1.01, 1.11)* | 1.03 (1.01, 1.06)* | 1.02 (1.00, 1.03)* | 1.03 (1.01, 1.05)** | 1.01 (0.99, 1.03) | -0.07 (-0.44, 0.30) |
| GrimAgeAA | -1.10 (-1.51, -0.70)*** | -0.14 (-0.25, -0.02)* | 1.11 (1.05, 1.17)*** | 1.07 (1.04, 1.10)*** | 1.03 (1.01, 1.04)*** | 1.04 (1.02, 1.07)** | 1.04 (1.02, 1.06)*** | 0.47 (0.05, 0.88)* |
| **Model 2** |  |  |  |  |  |  |  |  |
| HorvathAA | -0.06 (-0.30, 0.17) | 0.08 (0.02, 0.15)* | 0.98 (0.95, 1.02) | 0.99 (0.98, 1.03) | 1.00 (0.99, 1.01) | 0.99 (0.98, 1.01) | 1.00 (0.99, 1.01) | -0.10 (-0.36, 0.16) |
| HannumAA | 0.00 (-0.24, 0.24) | 0.05 (-0.02, 0.12) | 0.96 (0.93, 1.00)* | 0.99 (0.97, 1.01) | 1.00 (0.99, 1.00) | 0.99 (0.98, 1.01) | 1.00 (0.99, 1.01) | -0.12 (-0.38, 0.14) |
| PhenoAgeAA | -0.29 (-0.65, 0.07) | 0.02 (-0.08, 0.12) | 1.04 (0.99, 1.09) | 1.02 (0.99, 1.05) | 1.01 (1.00, 1.02) | 1.03 (1.01, 1.05)* | 1.00 (0.99, 1.02) | -0.11 (-0.51, 0.27) |
| GrimAgeAA | -0.95 (-1.45, -0.44)*** | -0.08 (-0.23, 0.06) | 1.10 (1.02, 1.19)** | 1.06 (1.01, 1.10)** | 1.02 (1.00, 1.03)* | 1.02 (0.99, 1.05) | 1.03 (1.00, 1.05) | 0.11 (-0.45, 0.67) |

Model 1: adjusted for age and sex

Model 2: Model 1 + life course social class trajectory, smoking, physical activity, BMI

***significant at the p<.001 level; ** significant at the p<.01 level * significant at the p<.05 level
